# Patient-specific functional genomics and disease modeling suggest a role for LRP2 in hypoplastic left heart syndrome

**DOI:** 10.1101/835439

**Authors:** Jeanne L. Theis, Georg Vogler, Maria A. Missinato, Xing Li, Almudena Martinez-Fernandez, Tanja Nielsen, Stanley M. Walls, Anais Kervadec, Xin-Xin I Zeng, James N. Kezos, Katja Birker, Jared M. Evans, Megan M. O’Byrne, Zachary C. Fogarty, André Terzic, Paul Grossfeld, Karen Ocorr, Timothy J. Nelson, Timothy M. Olson, Alexandre R. Colas, Rolf Bodmer

**Author notes:** To whom correspondence should be addressed: Rolf Bodmer, PhD, Development, Aging and Regeneration Program, Sanford Burnham Prebys Medical Discovery Institute,10901 N Torrey Pines Rd., La Jolla, CA92037, USA. Equal contributions (Jeanne L. Theis, Georg Vogler, Maria A. Missinato). Equal contributions (Timothy J. Nelson, Timothy M. Olson, Alexandre R. Colas, Rolf Bodmer).

## Abstract

Congenital heart diseases (CHD), such as hypoplastic left heart syndrome (HLHS), are considered to have complex genetic underpinnings that are poorly understood. Here, an integrated multi-disciplinary approach was applied to identify novel genes and underlying mechanisms associated with HLHS. A family-based strategy was employed that coupled whole genome sequencing (WGS) with RNA sequencing of patient-derived induced pluripotent stem cells (iPSCs) from a sporadic HLHS proband-parent trio to identify, prioritize and functionally evaluate candidate genes in model systems. Consistent with the hypoplastic phenotype, the proband’s iPSCs had reduced proliferation capacity. Filtering WGS for rare de novo, recessive, and loss-of-function variants revealed 10 candidate genes with recessive variants and altered expression compared to the parents’ iPSCs. siRNA/RNAi-mediated knockdown in generic human iPSC-derived cardiac progenitors and in the *in vivo Drosophila* heart model revealed that LDL receptor related protein *LRP2* and apolipoprotein *APOB* are required for robust hiPSC-derived cardiomyocyte proliferation and normal hear structure and function, possibly involving an oligogenic mechanism via growth-promoting WNT and SHH signaling. *LRP2* was further validated as a CHD gene in a zebrafish heart model and rare variant burden testing in an HLHS cohort. Collectively, this cross-functional genetic approach to complex congenital heart disease revealed LRP2 dysfunction as a likely novel genetic driver of HLHS, and hereby established a scalable approach to decipher the oligogenic underpinnings of maladaptive left heart development.

**One sentence summary:** Whole genome sequencing and a multi-model system candidate gene validation - human iPSC-derived cardiomyocytes and *Drosophila* and zebrafish hearts - identified lipoprotein *LRP2* as a new potential driver in congenital heart disease and suggests a deficit in proliferation as a hallmark of hypoplastic left heart syndrome.

## INTRODUCTION

Hypoplastic left heart syndrome (HLHS) is a congenital heart disease (CHD) characterized by underdevelopment of the left ventricle, mitral and aortic valves, and aortic arch. Variable phenotypic manifestations and familial inheritance patterns, together with the numerous studies linking it to a diverse array of genes, suggest that HLHS is genetically heterogeneous and may have significant environmental contributors (*1–4*). In this scenario, synergistic combinations of filtering and validating approaches are necessary to prioritize candidate genes and gene variants that may affect cardiogenic pathways throughout the dynamic process of human heart development.

Although the cellular mechanisms for HLHS remain poorly characterized, a recent study reported generation of the first animal model of HLHS. Based on a digenic mechanism, mice deficient for HDAC-associated protein-encoding *Sap130* and protocadherin-coding *Pcdha9* exhibited left ventricular (LV) hypoplasia that was likely due – at least in part – to defective cardiomyocyte proliferation and differentiation, and increased cell death (*5*). Similarly in humans, Gaber et al. (*6*) provide evidence that HLHS-LV samples have more DNA damage and senescence with cell cycle arrest, and fewer cardiac progenitors and myocytes than controls. These observations suggest that impaired cardiomyocyte proliferation could be a mechanism contributing to HLHS pathogenesis, although pathogenic genes controlling this process in humans remain to be identified and validated. Therefore, new synergistic experimental approaches that functionally validate cellular mechanisms of defective cardiogenesis are needed to probe the oligogenic hypothesis of left-sided heart defects, such as HLHS (*7, 8*).

Over the last decade, induced pluripotent stem cells (iPSCs) have provided a revolutionary experimental tool to reveal aspects of the cellular manifestations associated with disease pathogenesis (*9–11*). Progress in next-generation sequencing technology allows rapid whole genome DNA and RNA sequencing, thereby providing access to high-resolution and personalized genetic information. However, the interpretation of patient-specific sequence variants is often challenged by uncertainty in establishing a pathogenic link between biologically relevant variant(s) and a complex disease (*12*).

Testing numerous potential human disease-susceptibility genes in a mammalian *in vivo* model has been challenging because of high costs and low throughput. *Drosophila* with its genetic tools has emerged as the low cost, high throughput model of choice for human candidate disease gene testing, including neurological and cardiac diseases(*13–15*). *Drosophila* has been established as an efficient model system to identify key genes and mechanisms critical for heart development and function that served as prototypes for vertebrate/mammalian studies, including in zebrafish and mice, due to high degree of conservation of genetic pathways and reduced genetic complexity (*16*), e.g the first cardiogenic transcription factor *Nkx2-5/tinman,* discovered in *Drosophila* (*17, 18*), marked the beginning of a molecular-genetic understanding of cardiac specification (*19–25*).

For this study, we combined whole-genome sequencing (WGS), iPSC technology and model system validation with a family-based approach to identify and characterize novel HLHS-associated genes and mechanisms. This approach led us to identify LRP2 as a major regulator of cardiac cardiomyocyte proliferation of hiPSCs and heart development and maturation in both *Drosophila* and zebrafish. Consistent with our model system findings, burden analysis revealed that rare and predicted deleterious missense *LRP2* variants were enriched in HLHS patients as compared to healthy controls. Finally, we found evidence that suggests *LRP2* regulates cardiac proliferation and differentiation by modulating growth-associated WNT, SHH and TP53 pathways. We conclude there may be a novel genetic link between *LRP2* function and cardiac growth in the etiology of HLHS. Furthermore, our integrated multidisciplinary high-throughput approach establishes a scalable and synergistic gene discovery platform in genetically complex forms of human heart diseases.

## RESULTS

### Transcriptome and cell cycle activity are altered in HLHS patient-derived iPSCs and CMs

This study analyzed a family comprised of unrelated parents and their three offspring (‘5H’ family; Figure **1A**). The male proband (II.3) was diagnosed with non-syndromic HLHS by physical examination and echocardiography, which demonstrated aortic and mitral valve atresia, virtual absence of the left ventricular cavity, and severe aortic arch hypoplasia. He was born prematurely at 29 weeks gestation and underwent staged surgical palliation at 2 and 11 months of age. Conversion to a fenestrated Fontan circulation at 3 years of age failed owing to systolic and diastolic heart failure, necessitating early take-down. The patient subsequently died of multi-organ system failure. Echocardiography revealed structurally and functionally normal hearts in the proband’s mother (I.2), father (I.1) and siblings (II.1 and II.2). Maternal history is notable for 4 miscarriages.

**Figure 1:**
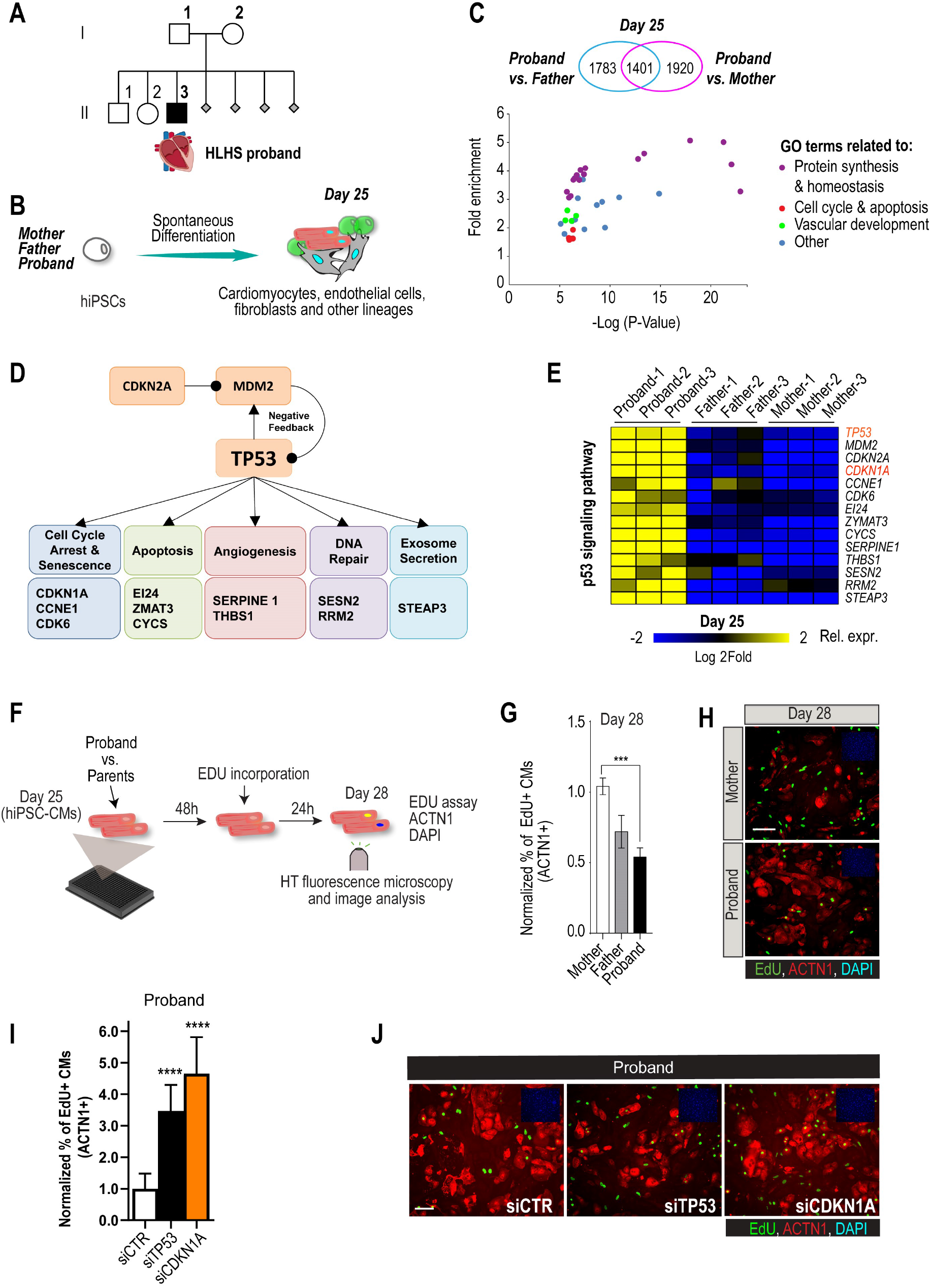
Family-based iPSC characterization for HLHS. (**A**) Pedigree of family 5H: proband with HLHS (black symbol), relatives without CHD (white symbols), miscarriages (gray diamonds). (**B**) Schematic for family-based iPSC production and characterization. (**C**) Whole genome RNA sequencing identified 1401 concordantly DETs between proband and parents. (**D**) KEGG pathway analysis shows enrichment of DETs in TP53 pathway. (**E**) Heatmap of p53 signaling pathway associated genes in probands vs parents. (**F**) Schematic describing EdU incorporation assay in hiPSC-CMs. 5000 cells/well were plated in 384 well plate. After 48 hours EdU was added to the media and left incorporate for 24 hours. Cells were then fixed and stained (**G**) Graph representing quantification of EdU+ cardiomyocytes in HLHS 5H family-derived iPSC-CMs. ***p<0.001 One-way ANOVA. (**H**) Representative images of iPSC-CMs derived from mother (Top) and proband (Bottom), stained for EdU, ACTN1 and DAPI. Scale bar: 50 μm. (**I**) Quantification of EdU incorporation assay in 5H proband iPSC-CM upon KD of TP53 or CDKN1A. ****p<0.0001, One-way ANOVA. (**J**) Representative images of 5H proband iPSC-CM stained for EdU and ACTN1 upon KD of TP53 or CDKN1A at day 28. Scale bar: 50 μm.

Patient-derived iPSCs are a valuable tool to investigate heart defects, such as those observed in HLHS (*3, 26*). In this study, iPSCs from the mother (I.2), father (I.1) and HLHS proband (II.3) were generated (*27*) to investigate differences in transcriptional profiles potentially associated with HLHS, cells from the proband-parent trio were differentiated to day 25, using a cardiogenic differentiation protocol and processed for subsequent RNA sequencing (Figure **1B**). In this *in vitro* cellular context, bioinformatic analysis revealed that 5,104 differentially expressed transcripts (DETs) in D25 differentiated samples between proband vs. mother/father (Supplementary Table 1, Benjamini-corrected p<0.001). We found that 1,401 DETs were concordantly differentially expressed between proband and both parents (Figure **1C,** Supplementary Figure **1A-C**, Supplementary Tables **1,2**). Consistent with previous observations in HLHS fetuses (*6*), KEGG analysis revealed TP53 pathway enrichment (Figure **1D**), including cell cycle inhibition (Figure **1E** and Supplementary Figure **1A-C**), suggesting that cell proliferation may be affected in proband cells.

To test this hypothesis, we measured cell cycle activity in proband and parent hiPSC-derived cardiomyocytes (hiPSC-CMs) using an EdU-incorporation assay (Figure **1F**). Indeed, proband hiPSC-CMs exhibited reduced percentage of EdU-positive cells as compared to parents (Figure **1G,H**). To further evaluate whether a potentially reduced proliferative activity is a more general phenotypic hallmark of HLHS cells, we evaluated the proliferative status of two additional HLHS family trios that were available to us from the HLHS cohort at Mayo Clinic (‘75H’, ‘151H’). Consistent with our findings with 5H family trio cells (Figure **1G,H**), the proband cells of families 75H and 151H also exhibited significant reduction of proliferative activity as compared to the parents using the EdU-incorporation assay (Supplementary Figure **1D-E**). Given the upregulation of potent cell cycle inhibitors *TP53* or *CDKN1A* in 5H proband cells (Figure **1E**), we tested whether impaired proliferation could be caused by the observed elevated *TP53* and/or *CDKN1A* mRNA levels. Indeed, siRNA-mediated knockdown (KD) of TP53 and CDKN1A KD in proband hiPSC-CMs significantly increased EDU incorporation as compared to siControl (Figure **1I,J**). These findings are consistent with a causal connection between HLHS and a TP53/CDKN1A-dependent blockade of CM proliferation, and is consistent with observations made in both HLHS fetuses (*6*) and a HLHS mouse model (*5*).

### Family-based WGS, variant filtering, and transcriptional profiling identified 10 candidates

Array comparative genome hybridization ruled out a chromosomal deletion or duplication in the proband. WGS was performed to identify potentially pathogenic coding or regulatory single nucleotide variants (SNVs) or insertion/deletions (INDELs). First, we ruled out pathogenic variants within 42 genes comprising a congenital heart disease genetic testing panel (Invitae, San Francisco, CA). To identify novel HLHS candidate genes, WGS of the family quintet was filtered for rare *de novo,* recessive and loss-of-function variants with predicted impact on protein structure or expression, yielding 114 variants in 61 genes (Supplementary Figure **2**, Supplementary Table **3**). We next prioritized genes most likely to drive downstream pathways of dysregulated cardiogenesis in the HLHS proband by cross-referencing these candidate genes with 3,816 DETs identified in undifferentiated iPSC at day 0 (d0) (Supplementary Table **4**) and 5,104 DETs identified at day 25 (d25) differentiated cell lineages (Supplementary Table **1**). Ten genes harboring compound heterozygous (*7*), hemizygous (*2*), or homozygous (*1*) recessive variants (Table **1**), absent in the unaffected siblings, were found to be differentially expressed within the HLHS proband’s iPSCs at d0 and d25: *HSPG2, APOB, LRP2, PRTG, SLC9A1, SDHD, JPT1, ELF4, HS6ST2* and *SIK1* (Figure **2A**).

**Figure 2:**
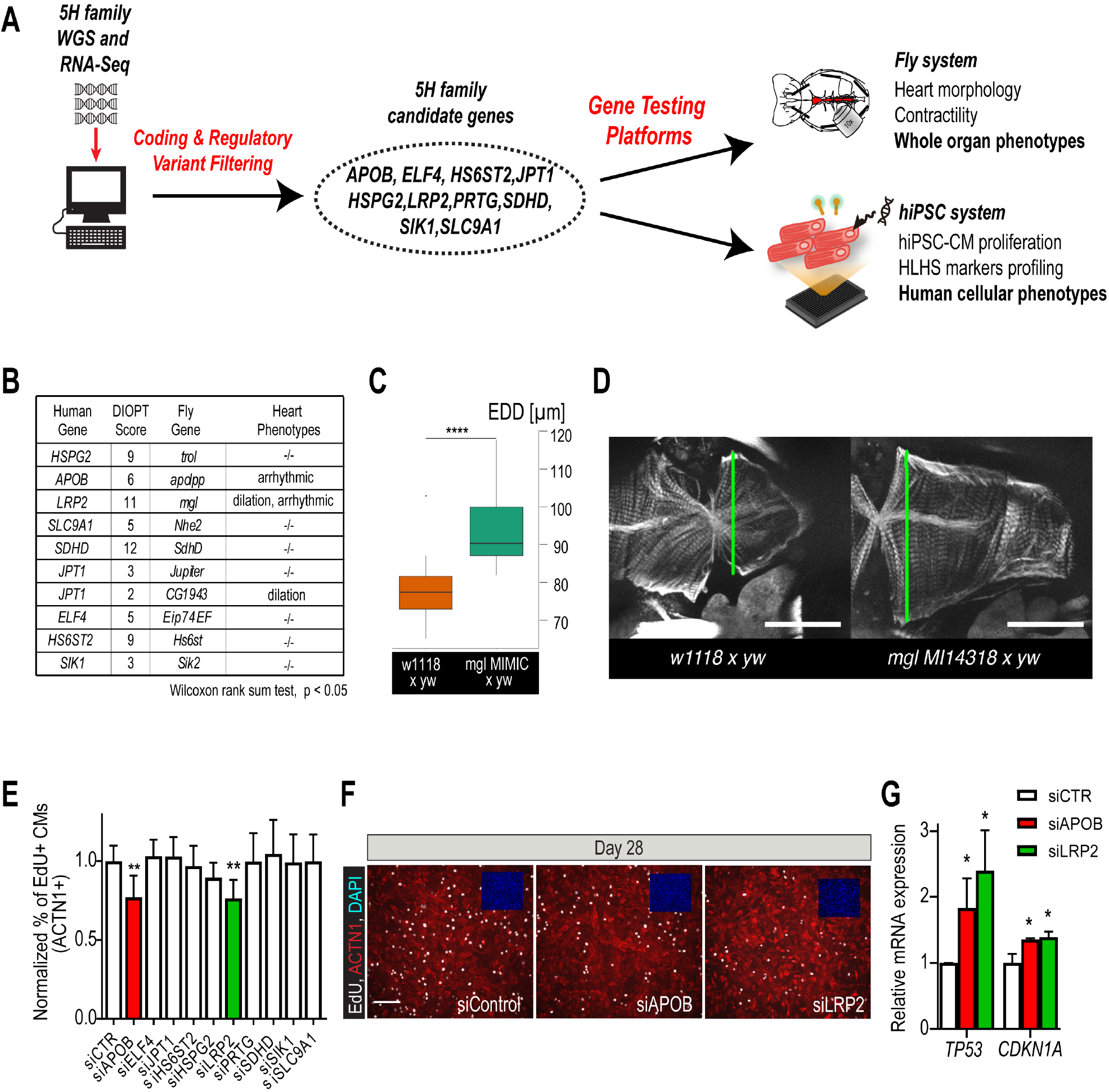
Whole genome and RNA sequencing identify HLHS candidate genes. (**A**) An iterative, family-based variant filtering approach based on rarity, functional impact, and mode of inheritance and RNA sequencing data were used to filter for transcriptional differences yielding 10 candidate genes. Candidate genes were further tested in hiPSC-CM and *in-vivo* model. (**B**) Human candidate genes and corresponding *Drosophila* ortholog as determined by DIOPT score (*confidence score: number of databases reporting orthology). Listed are heart phenotypes upon gene candidate loss of function. (**C,D**) Example of fly hearts heterozygous for *LRP2/mgl* show increased end-diastolic diameters (EDD, measured at green line in **D**). Wilcoxon rank sum test: ***p<0.001. (**E**) Graph representing EdU incorporation assay results of candidate gene KD in hiPSC-CM. KD of APOB (red bar) or LRP2 (green bar) reduced EdU incorporation. **p<0.01 One-way ANOVA. (**F**) Representative images of hiPSC-CMs stained for EdU, ACTN1 and DAPI. Scale bar: 50 μm. (**G**) qPCR results of TP53 and CDKN1A in hiPSC-CM upon KD of APOB or LRP2. *p<0.05 One-way ANOVA.

**Table 1.** Recessive Variants Identified in 10 Candidate Genes

| Gene | Mode of Inheritance | Functional Impact | Transcript Variant | Protein Variant | Inheritance | Genotype in Brother (II.1) | Genotype in Sister (II.2) | gnomAD* MAF (%) | dbSNP ID |
| --- | --- | --- | --- | --- | --- | --- | --- | --- | --- |
| *HSPG2* | Cmpd Het | missense | c.2074G>A;<br>c.2077G>A | p.V692M;<br>p.V693M | Maternal | WT | WT | 0.288 | 143669458 |
|  |  | missense | c.326G>A | p.R109Q | Paternal | Het | Het | 0 | 773796176 |
|  |  | promoter | c.-227C>A |  | Paternal | WT | WT | 1.392 | 566166086 |
| *SLC9A1* | Cmpd Het | promoter | c.-906T>C |  | Paternal | Het | Het | 1.227 | 114101904 |
|  |  | promoter | c.-947T>G |  | Maternal | WT | WT | 27.175 | 11588974 |
|  |  | promoter | c.-1085A>G |  | Paternal | Het | Het | 0.841 | 116299278 |
|  |  | ENCODE TFBS | c.-1138C>T |  | Paternal | Het | Het | 0.93 | 75089536 |
|  |  | promoter | c.-1311G>A |  | Paternal | Het | Het | 0.93 | 77414471 |
| *APOB* | Cmpd Het | missense | c.13441G>A | p.A4481T | Maternal | WT | Het | 2.475 | 1801695 |
|  |  | missense | c.751G>A | p.A251T | Paternal | Het | WT | 0.071 | 61741625 |
| *LRP2* | Cmpd Het | missense | c.9613A>G | p.N3205D | Maternal | WT | WT | 0.407 | 35734447 |
|  |  | missense | c.170C>T | p.A57V | Paternal | WT | WT | 0.032 | 115350461 |
| *SDHD* | Cmpd Het | promoter | c.-815G>C;<br>c.129+547C>G |  | Maternal | Het | WT | 0.573 | 117661257 |
|  |  | ENCODE TFBS | c.-205G>A;<br>c.66C>T | p.A22A | Paternal | WT | WT | 0.241 | 61734353 |
|  |  | missense | c.34G>A; c.-173C>T | p.G12S | Maternal | Het | WT | 0.729 | 34677591 |
| *PRTG* | Cmpd Het | microRNA Binding Site | c.*3501T>G |  | Paternal | Het | WT | 0.739 | 77181316 |
|  |  | microRNA Binding Site | c.*2678A>G |  | Maternal | WT | Het | 0.019 | 756136447 |
| *HN1* | Cmpd Het | ENCODE TFBS | c.56+617C>T;<br>c.-903C>T; c.-178+617C>T;<br>c.-590C>T |  | Maternal | Het | WT | 3.764 | 117213586 |
|  |  | promoter | c.-1748A>C;<br>c.-719A>C; c.-486A>C |  | Paternal | WT | Het | 0.816 | 73995795 |
| *SIK1* | Hom Rec | missense | c.2087C>T | p.P696L | Maternal and Paternal | WT | Het |  | 1256991707 |
| *ELF4* | X-Linked | missense | c.1144G>A | p.V382I | Maternal | WT | Het | 0.025 | 148953158 |
| *HS6ST2* | X-Linked | missense | c.948-40041G>A;<br>c.1046G>A | p.R349Q | Maternal | WT | Het | 0.146 | 201239951 |

3 Cmpd Het, compound heterozygous; Het, heterozygous; Hom Rec, homozygous 4 recessive; MAF, minor allele frequency; WT, wild type.
\*At study initiation the ESP database was used to set the 3% allele frequency filter. Updated frequencies are shown based on the newer gnomAD database curation which would now eliminate *SLC9A1* and *HN1* as candidate genes.

### Knockdown of candidate gene orthologs in *Drosophila* heart

In order to determine whether these variants occurred within genes that could be important for cardiac differentiation *in vivo,* we took advantage of our established *Drosophila* heart development model and functional analysis tools (Supplementary Figure **3A**) (*14*). We hypothesized that genes critical for the *Drosophila* heart have conserved roles also in humans, as previously observed (*19, 20, 28*). Predicted by DIOPT database (*29*) to have orthologs in *Drosophila* (Figure **2B**), we analyzed 9 genes using heart-specific RNAi KD. By *in vivo* heart structure and function analysis (*13*), we found that KD of *LRP2 (mgl)* and *JPT1 (CG1943)* caused dilated heart phenotypes, while KD of *APOB (apolpp),* a circulating lipoprotein ligand, and again *LRP2 (mgl)* also resulted in arrhythmias (Figure **2B-D**, Supplementary Figure **3** and Supplementary movies **S1**, **S2**, **S3**), suggesting developmental or remodeling defects of cardiac structure and function.

Since HLHS is likely oligogenic (*30, 31*), functional requirements for some genes involved in HLHS might only become apparent in combination with variants in other cardiac-relevant genes. To test this, we examined the candidates also in the heterozygous background for *tinman/NKX2-5,* which in humans is well-known to contribute to a variety of CHD/HLHS manifestations.(*1, 26, 32, 33)* In this *in vivo* context, heart-specific KD of 2 out of 9 genes, namely *HSPG2/Perlecan (trol),* involved in extracellular matrix assembly (*34*), and Succinate dehydrogenase subunit D *SDHD (SdhD)* exhibited a constricted phenotype (Supplementary Figure **3G,H**). These findings demonstrate that our bioinformatic candidate gene prioritization identified several conserved candidates as cardiac relevant.

### *LRP2* and *APOB* regulate proliferation in human iPSC-derived cardiomyocytes

Decreased proliferation of left ventricular cardiomyocytes is emerging as a phenotypic hallmark of HLHS (*5, 6*) (see also Figure **1G,H** and Supplementary Figure **1D**), suggesting that cell cycle impairment may an important contributor to the disease. Thus, we asked whether siRNA-mediated KD of the prioritized 10 candidate genes from the 5H family trio affects proliferation of generic hiPSC-CM (*35*). Remarkably, two of the genes causing cardiac abnormalities when knocked down in *Drosophila, LRP2* and *APOB,* also caused a marked reduction of EdU+ hiPSC-CMs (ACTN1+) and overall hiPSC cell numbers (Figure **2E,F** and Supplementary Figure **4A,B**. Notably, we also observed an upregulation of cell cycle inhibitors and apoptosis genes (Supplementary Figure **4C**), including TP53 and CDKN1A (Figure **2G**), as well as a downregulation of cell cycle genes (Supplementary Figure **4B,C**). Collectively, these data identify *LRP2* and *APOB* as regulators of cell cycle and apoptosis in hiPSC-CMs, thus potentially contributing to the developmental cardiac impairment in HLHS patients.

### Rare variant analysis in HLHS cohort reveals enrichment in *LRP2*

In order to determine disease relevance of candidate genes functionally validated in both systems, we asked whether the frequency of rare and predicted-damaging variants in *LRP2* and *APOB* would be higher in a cohort of 130 HLHS cases compared to 861 control individuals. Remarkably, HLHS patients had a ~3-fold increase in the frequency of rare, predicted-damaging *LRP2* missense variants compared to healthy controls (10% versus 3.4%; p=0.0008)(Figure **3A;** Supplementary Table **6**). Aamong the 13 patients who carried a *LRP2* variant (Figure **3A,B**), 3 shared the same variant (N3205D) with the 5H proband (Figure **3B**) (Supplementary Table **7**). Of note, 13 of the 130 HLHS cases (including the index family proband) possessed <80% of ancestral Caucasian alleles, while all controls possessed ≥80%. Four of the 13 cases had rare, predicted-damaging missense variants in *LRP2* however all variants assessed were required to be rare in all racial populations. To eliminate the potentially confounding variable of race a Caucasian-only sub-analysis was performed, resulting in a less significant p-value for rare, predicted-damaging missense variants (7.7% versus 3.4%; p=0.05). However, removal of the predicted-damaging (CADD) restriction on rare *LRP2* variants among Caucasians revealed significant enrichment in cases (p=0.0035), most notably in missense and intronic variants (p=0.0178 and 0.0082, respectively, Supplementary Table **8**). Population-based allele frequencies, CADD scores, and location of variants within functional protein binding domains, active histone marks, or transcription factor binding sites was not different between cases and controls.

**Figure 3:**
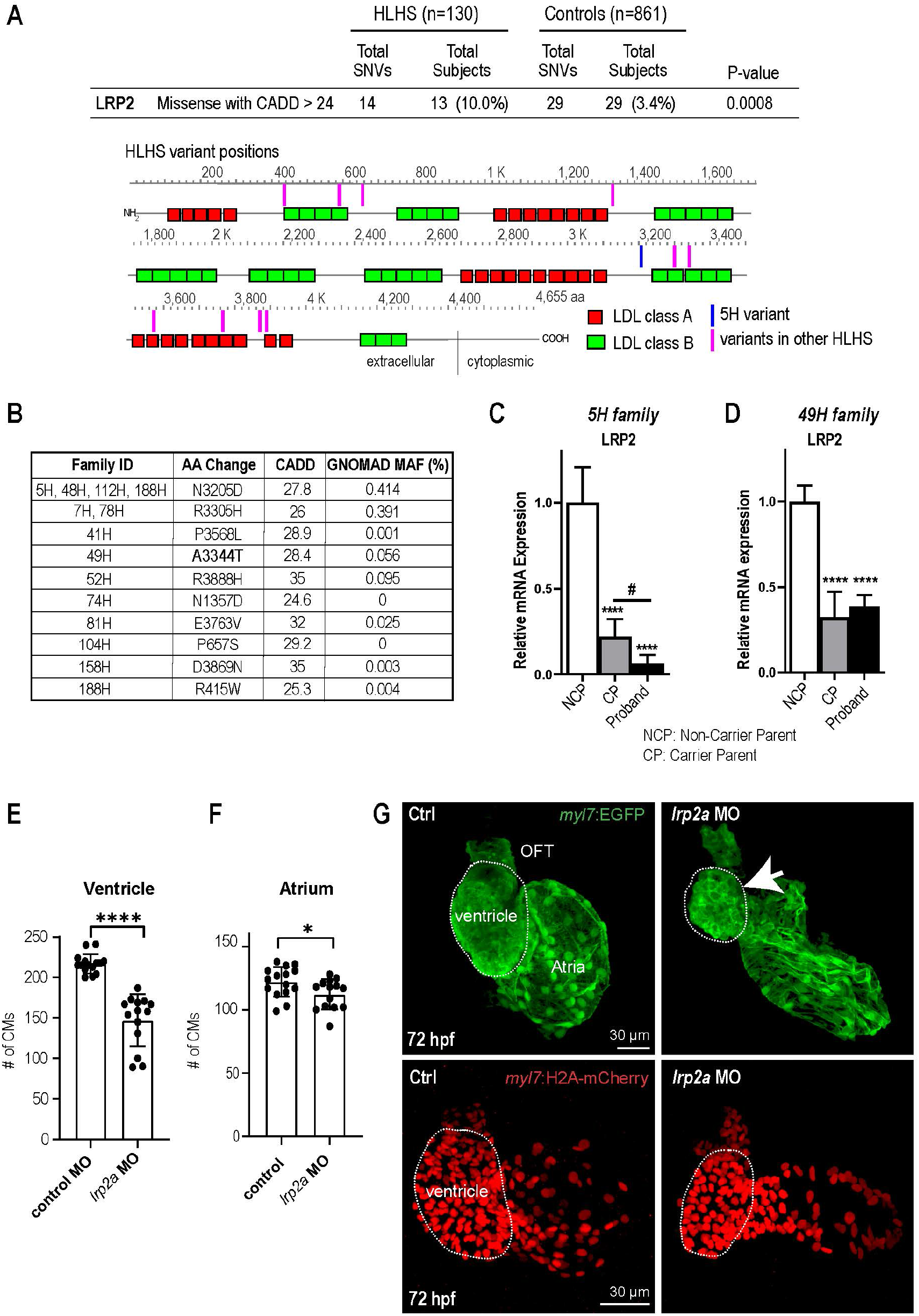
Identification of LRP2 as potential HLHS-susceptibility gene. (**A**) Cohort-wide analysis of LRP2 variants shows significant enrichment for SNVs in HLHS patients compared to control populations. Variants (blue/magenta) are found throughout LRP2 protein. (**B**) Table listing the HLHS families carrying LRP2 variants. (**C,D**) qPCR of LRP2 in 5H family (**C**) and in 49H family (**D**) showing LRP2 downregulation in carrier parent and proband compared to the non-carrier parent. ****p<0.0001 One-way ANOVA #p<0.05 One-way ANOVA. (**E**) Cardiomyocyte count in zebrafish morphants at 72 hpf were significantly reduced in the ventricle. (**F**) Atrial cardiomyocyte number was also reduced in morphants but to a lesser extent than in ventricles. *p<0.05; ****p<0.0001 Unpaired two-tail student t-test. (**G**, top panel) Embryonic fish hearts were visualized by EGFP expression in the *myl7:EGFP* transgenic background (green) at 72 hpf. *lrp2a* morphants hearts were dysmorphic and much smaller (arrow) compared to controls. Dotted circles outline the ventricles. (**G**, lower panel) *myl7:H2A-mCherry* transgenic background identifies cardiomyocyte nuclei used for quantifying cardiomyocytes during development in E, and F. Dotted trace outlines the ventricle. Scale bars: 30 μm.

In a next step, we sought to determine whether LRP2 levels might be affected in probands with rare, predicted-damaging variants in LRP2 coding sequence. We profiled *LRP2* transcripts levels in patient-derived iPSCs of the 5H family as well as another family, 49H, both harboring heterozygous variants with a CADD score above 24 (Figure **3B**), inherited from one of the parents. Interestingly, qPCR results showed that LRP2 mRNA levels were lower in the probands of both families, as well as in the parent carrying the variant (CP), compared to non-variant carrying parent (NCP) (Figure **3C,D**). This corroborates the idea that these LRP2 variants (N3205D and A3344T) may be causing a genetic loss-of-*LRP2*-function in the 5H and 49H families, thus potentially contributing to the proband’s HLHS. However, there are likely other contributing factors besides the presence of the *LRP2* variants, since echocardiography excluded CHD in carrier parents.

### Loss-of-function in zebrafish *LRP2* causes a hypoplastic ventricular phenotype

In order to evaluate the role of LRP2 during heart development in a vertebrate model, we injected a morpholino as well as sgRNA/CRISPR directed against *LRP2 (lrp2a)* in zebrafish embryos and evaluated the effect on heart morphology and function at 72 hpf. Overall body morphology was similar for morphant and F0 CRISPR edited larva at 72 hpf, compared to controls (Supplemental Figure **5A-C**). However, hearts from larva with reduced *lrp2a* function displayed a hypoplastic phenotype with decreased CM number (Figure **3E-G**) and dose-dependent reductions in ventricular chamber dimensions in morphants (Supplemental Figure **5D,E**). Loss-of-*lrp2a*-function also compromised ventricular contractility and caused bradycardia in both morphants and CRISPR edited larva (Supplementary Figure **5F,G**). Collectively, our data suggest that *LRP2* plays a crucial role during heart development by regulating cardiomyocyte generation most prominently in the ventricular chamber.

### Potential regulatory network of validated gene candidates

In order to delineate how 5H family candidate genes testing positive in one or both of the our validation systems, *APOB, HS6ST2, HSPG2, JPT1, LRP2,* might affect signaling homeostasis in proband cells, we assembled a gene network containing these 5 genes and their first neighbors (as genetic and protein-protein interactions, BioGRID) (Figure **4A**; Supplementary Table **5**). Strikingly, in addition to TP53 pathway misregulation (Figure **1D-E**, Figure **2G** and Supplementary Figure **4**) all 5 genes were connected to WNT and SHH signaling cascades, both key regulators of cardiac differentiation and proliferation (*36, 37*) (Figure **4A**). Notably, RNA-seq analysis of the proband cells confirmed this network prediction: the negative regulator of SHH pathway, *PTCH1*, was upregulated, while agonists of WNT signaling pathway, *WNT1/3a/8a/10b* and *FZD10* (*38*), were downregulated, compared to parental cells, suggesting these pathways are attenuated in the proband.

**Figure 4:**
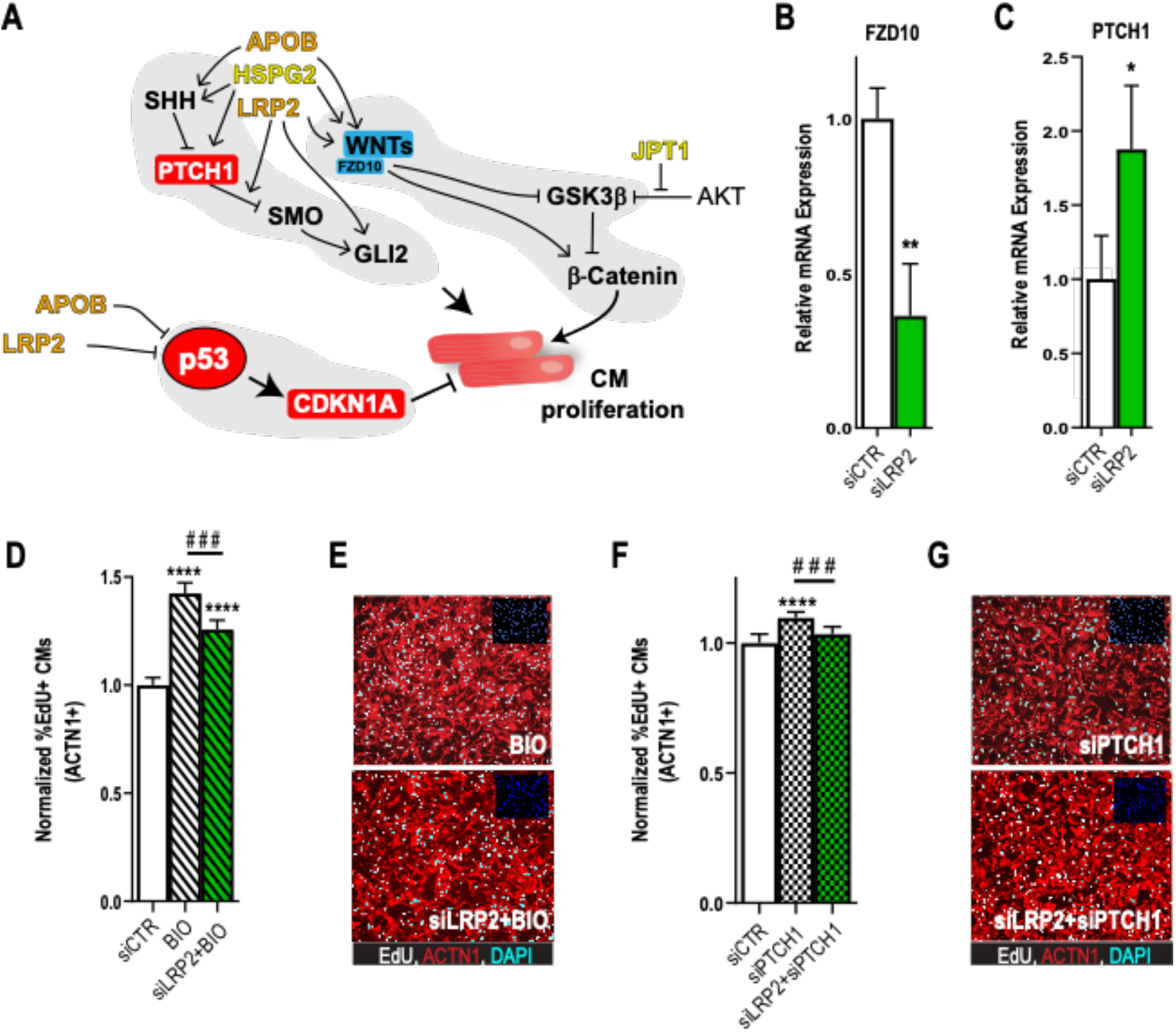
Potential role for SHH, WNT and LRP2 in HLHS. (**A**) A gene network integrating family-centric HLHS candidate genes with heart development. ORANGE – genes with cardiac phenotypes in iPSC/*Drosophila* assays. YELLOW – other candidate genes with *Drosophila* phenotypes. RED – Genes up-regulated in proband iPSCs vs. parents. BLUE – Genes downregulated down-regulated in proband iPSCs vs. parents. (**B**,**C**) qPCR for FZD10, a WNT-pathway associated gene, (**B**) and for PTCH1, a SHH pathway associated gene (**C**) upon LRP2 KD in hiPSC-CM. **p<0.01, *p<0.05 Student’s t-test. (**D**) Quantification of EdU incorporation assay in hiPSC-CM upon LRP2 KD in combination with BIO, a WNT inhibitor. ***or###p<0.001, ****p<0.0001, One-way ANOVA. (**E**) Representative images of hiPSC-CM stained for EdU and ACTN1. Scale bars: 50 μm. (**F**) Quantification of EdU incorporation assay in hiPSC-CM upon LRP2 in combination with PTCH1 KD, a SHH associated gene. ***p<0.001, ****p<0.0001, Oneway ANOVA. (**G**) Representative images of hiPSC-CM stained for EdU and ACTN1. Scale bars: 50 μm.

To further examine a potential link between the 5H proband vs. parental expression profiles and LRP2, we asked whether LRP2 could regulate WNT- and/or SHH-associated genes. Consistent with WNT/SHH-centric network predictions and 5H family RNA-seq data, KD of LRP2 led to a reduction in *FZD10* expression concomitant with increased *PTCH1* RNA levels (Figure **4B,C**), although WNT1/3a/8/10a were not affected. Given that LRP2 regulates key component of WNT *(FZD10)* and SHH *(PTCH1)* pathways, we next asked whether LRP2 function was required for WNT and/or SHH-induction of CM proliferation. For this purpose, we used the WNT agonist BIO (*39*) and siRNA against PTCH1 (*40*). Strikingly, LRP2 KD significantly reduced both BIO- and siPTCH1-induced proliferation in hiPSC-CMs (Figure **4D-G**), suggesting that LRP2 is required for both WNT- and SHH-regulated cardiomyocyte proliferation.

## DISCUSSION

### Integrated multidisciplinary disease gene discovery and modeling

Unraveling the molecular-genetic etiology of HLHS pathogenesis is crucial for (*1*) our ability to provide reliable prenatal diagnostics to families and (*2*) to develop novel approaches to treat the disease. As an important step towards these goals, our integrated multidisciplinary approach identifies rare variants in *LRP2* as a molecular signature enriched in HLHS patients, consistent with our mechanistic analysis in model systems.

For this study, we used a powerful combination of high-throughput DNA/RNA patient sequencing coupled with high-throughput functional screening in model systems enabling to probe gene function (alone or in combination) on a wide array of cellular processes that are deployed during heart formation. For validation in model systems, we have established an integrated multi-site and multidisciplinary pipeline that systematically evaluates the functional role of genes presenting rare and deleterious variants in HLHS patients in hiPSC and *Drosophila* and zebrafish heart models. As a main objective – identify genes and potential mechanisms associated with HLHS – our study highlights rare, predicted-damaging *LRP2* missense variants as 3-fold enriched in 130 HLHS patients compared to 861 controls. Validation of this gene in hiPSC and the *Drosophila* and zebrafish heart models demonstrated a requirement in cardiac proliferation and differentiation, and notably, systemic KD in zebrafish resulted in primarily in ventricular cardiac but not obvious skeletal muscle defects (see also Supplementary Figure **5**). Mutations in *LRP2* have been previously associated with cardiac defects in mouse(*41*) and in Donnai-Barrow Syndrome in humans (*41, 42*). However, *LRP2* has not previously been linked to HLHS within curated bioinformatic networks.

One pre-requisite to reduce the knowledge gap between patient genomes and clinical phenotypes is to establish reliable/quantifiable phenotypic links between HLHS-associated genes and their role during normal cardiac development. Also, given that each large-scale genomic study to identify CHD-associated genes can generate hundreds of candidates, we reasoned that our cardiac phenotypical platform should enable high-throughput functional screening strategies to accommodate rapid testing of a large number of genes. Although overall heart structure in flies differs from that in vertebrates, the fundamental mechanisms of heart development and function are remarkably conserved, including a common transcriptional regulatory network (*19, 20*), and a shared protein composition (*43*) as well as electrical and metabolic properties (*14, 44, 45).* This ‘convergent biology’ approach validated *LRP2* (and *APOB)* as HLHS candidate genes in both the *in vitro* and *in vivo* cardiac model systems. Importantly, variants in *LRP2* were not only found to be enriched in a cohort of 130 HLHS family trios, but also produced a ventricular hypoplastic phenotype in zebrafish embryos upon loss-of-*lrp2a*-function. Therefore, for further mechanistic understanding of complex CHD characterized by oligogenic etiologies this dual/triple model system testing approach enables assessment of gene function combinatorically and in sensitized genetic backgrounds (e.g. *tinman*/NKX2-5; see Supplemental Figure **3**).

### 5H pathogenic pathway: a potential role for SHH, WNT, p53 and cell proliferation in HLHS

Our current understanding of the molecular genetic causes of HLHS is very limited, despite clear genetic origins of disease (*46*). Past research on HLHS has yielded very few high-confidence gene candidates that may contribute to HLHS: *NOTCH1, NKX2-5* and *MYH6* have been implicated (*1, 3, 4),* but they are also associated with other CHDs.

Heart development is a complex process that involves the interaction of many pathways and tissue interactions, and a large number of genes have been implicated in various types of CHDs (*47*). The postulated oligogenic nature of HLHS likely is the result of an unfavorable combination of disease genes, and such a combination of alleles in turn could affect several, successive steps of heart development. This makes it extremely difficult to model the disease by single gene mutations. Current hypotheses of the etiology of HLHS include changes in cell cycle progression of myocytes, as well as altered blood flow (“no flow – no grow”) as a consequence of defects of valves or the outflow tract.

Interestingly, the only mouse HLHS model to date, a digenic mutant for *Sap130* and *pcdha9* (*5*), has a penetrance of less than thirty percent, indicating a profound role for subtle differences between genetic backgrounds. This mouse model suggests a separate mechanism with pcdha9 affecting aortic growth, whereas *Sap130* can exert a more severe HLHS-like phenotype, which might reflect a modular etiology of HLHS that separates valve and ventricular defects.

The gene network analysis that we have conducted points to the possibility that several of the prioritized candidate genes identified in the index 5H patient can have a modulatory impact on WNT and/or SHH via a diverse set of mechanisms (*36*). There is evidence that the three of the candidates with fly heart phenotypes – *Trol/HSPG2, Mgl/LRP2* and *Apolpp/APOB* – can alter WNT and SHH signaling (*48, 49*), consistent with both pathways being attenuated in the proband iPSCs. We hypothesize that a collective of likely hypomorphic genetic variants affects heart development leading to HLHS. Impaired ventricular growth could in addition be caused by changes in the p53 pathway, and our analysis of iPSC-derived cardiomyocytes suggests that p53 indeed depends on LRP2 levels. Such a multi-hit model of HLHS caused by sub-threshold hypomorphic alleles represents an attractive explanation of the disease.

In summary, this integrated multidisciplinary strategy of functional genomics using patientspecific iPSC combined with *in vivo* and human cellular model systems of functional validation postulates novel genetic pathways and potential polygenic interactions underlying HLHS. These complementary approaches enabled the deconvolution of patient-specific, complex polygenic risk factors potentially underlying HLHS and establish the novel groundwork for definitive mechanistic studies of interacting risk factors that contribute to defective cardiac development and adverse outcomes. This scalable approach promises more efficient discovery of novel genes associated more broadly with HLHS as multiple HLHS families can now be multiplexed in future studies.

## MATERIALS AND METHODS

### Study subjects

Written informed consent was obtained for the index family and an HLHS cohort, under a research protocol approved by the Mayo Clinic Institutional Review Board. Cardiac anatomy was assessed by echocardiography. Candidate genes were identified and prioritized by WGS of genomic DNA and RNA sequencing of patient-specific iPSC and cardiomyocytes. For variant burden analysis, controls were obtained from the Mayo Clinic Center for Individualized Medicine’s Biobank. Methods for genomic analyses, RNA Sequencing, iPSC technology, bioinformatics and statistics are described in the Online Appendix. Data are available in NCBI SRA database (see below for SRA Accession IDs).

### *Drosophila* and zebrafish heart function studies

*Drosophila* orthologs were determined using the DIOPT database (*29*), and RNAi lines were obtained from the Vienna Resource Drosophila Center (VDRC) stock center and crossed to the heart-specific *Hand^4.2^-Gal4* driver alone or in combination with one copy of the *tinman* loss-of-function allele *tin^EC40^* (*18*). Fly hearts were filmed and analyzed according to standard protocol (*13*).

In zebrafish, gene expression was manipulated using standard microinjection of morpholino (MO) antisense oligonucleotides (*50*). In addition, we performed targeted mutagenesis using CRISPR/Cas9 genome editing (*51–53*), to create insertion/deletion (INDEL) mutations in the *lrp2a* gene (F0). Zebrafish were raised to 72 hours post fertilization (hpf), immobilized in low melt agarose and the hearts were filmed and analyzed as for *Drosophila* (*13*).

### hiPSC-CM, siRNA transfection, EdU assay, Immunostaining, qRT-PCR

hiPSC-derived cardiomyocytes were produced as previously described (*35, 54*). Cardiomyocytes were plated in 384 wells and transfected with siRNAs (see Online Appendix). Two days after transfection, EdU was added to the media for 24 hours and fixed. EdU was detected using Click-it Plus EdU Imaging Kit (Life Technologies). For qRT-PCR experiments, total RNA was extracted using TRIzol and chloroform. 1ug of RNA was converted in cDNA using QuantiTect Reverse Transcription kit (Qiagen). Human primer sequences for qRT-PCR were obtained from Harward Primer Bank. At least 3 independent biological replicates were performed for each experiment.

### Statistical analysis

All statistical analysis for iPSC-derived cardiomyocytes were performed using GraphPad Prism version 8.0 (GraphPad Software, San Diego CA, USA). Statistical significance was analyzed by unpaired Student’s *t*-test, and one-way ANOVA and shown as mean ± SEM. P-values were considered significant when p < 0.05.

### Study Limitations

HLHS candidate gene selection was based on in silico predictive algorithms to filter for functional coding and regulatory variants. Our WGS filtering strategy, designed to identify majoreffect de novo, recessive and loss-of-function variants, did not include consideration of inherited, incompletely penetrant, autosomal dominant variants in other genes. The potential race-specific differences in *LRP2* variants require further study. Differential gene expression, which was functionally validated as a powerful filter for candidate variant prioritization, excluded functional variants that do not alter gene expression. The validating KD modeling systems are justified insofar as all 10 prioritized candidate genes harbored recessive alleles inherited from the proband’s unaffected parents, implicating a loss-of-function mechanism is likely in most cases. Not all human genes are conserved in *Drosophila,* but ~80% of disease-causing human genes have fly orthologs. While structural differences exist between hiPSC-CM, *Drosophila* and zebrafish hearts and human newborn cardiomyocytes, our combinatorial approach allows to uncover testable gene networks and interactions that is not feasible in mammalian model systems.

## Supporting information

Supplemental Movie S1 control fly heart

Supplemental Movie S2 APOB fly heart

Supplemental Movie S3 LRP2 fly heart

## Abbreviations

HLHS: hypoplastic left heart syndrome
CHD: congenital heart disease
WGS: whole-genome sequencing
SHH: sonic hedgehog
WNT: wingless/integrated

## Data availability

Sequencing data are deposited in the NCBI Sequence Read Archive (SRA) database with accession numbers: SRS1417684 (proband iPSCs), SRS1417685 (paternal iPSCs), SRS1417686 (maternal iPSCs), SRS1417695 (proband d25 differentiated cells), SRS1417696 (paternal d25 differentiated cells), SRS1417714 (maternal d25 differentiated cells).

## Acknowledgements

The authors acknowledge support from the Todd and Karen Wanek Family Program for Hypoplastic Left Heart Syndrome and the Medical Genome Facility Sequencing Core and Biobank within the Mayo Clinic Center for Individualized Medicine, Rochester, MN. We gratefully acknowledge the patient and family who participated in this study. We thank Sean Spearing, Prashila Amatya, Marco Tamayo and Bosco Trinh for excellent technical assistance.

## Sources of Funding

This work was supported by the Todd and Karen Wanek Family Program for Hypoplastic Left Heart Syndrome, Mayo Clinic Foundation, Rochester, MN (SAN-233970 to R.B. and A.R.C). This work was also supported by NIH (R01-HL054732 to RB).

## Competing interests

None.

## SUPPLEMENTARY MATERIAL

### SUPPLEMENTARY MATERIALS AND METHODS

#### Comparative genomic hybridization

To detect aneuploidy, array comparative genomic hybridization was performed using a custom 180K oligonucleotide microarray (Agilent, Santa Clara, CA), with a genome-wide functional resolution of approximately 100 kilobases. Deletions larger than 200 kilobases and duplications larger than 500 kilobases were considered clinically relevant.

#### Whole genome sequencing (WGS) and bioinformatics analyses of index family

Genomic DNA was isolated from peripheral blood white cells or saliva. WGS and variant call annotation were performed utilizing the Mayo Clinic Medical Genome Facility and Bioinformatics Core. Paired-end libraries were prepared using the TruSeq DNA v1 sample prep kit following the manufacturer’s protocol (Illumina, San Diego, CA). Each whole genome library was loaded into 4 lanes of a flow cell and 101 base pair paired-end sequencing was carried out on Illumina’s HiSeq 2000 platform using TruSeq SBS sequencing kit version 3 and HiSeq data collection version 1.4.8 software. Reads were aligned to the hg19 reference genome using Novoalign version 2.08 (http://novocraft.com) and duplicate reads were marked using Picard (http://picard.sourceforge.net). Local realignment of INDELs and base quality score recalibration were then performed using the Genome Analysis Toolkit version 1.6-9 (GATK).^1^ SNVs and INDELs were called across all samples simultaneously using GATK’s Unified Genotype with variant quality score recalibration (VQSR).^2^

Variant call format (VCF) files with SNV and INDEL calls from each family member were uploaded and analyzed using Ingenuity^®^ Variant Analysis™ software (QIAGEN, Redwood City, CA) where variants were functionally annotated and filtered by an iterative process. First, rare, high quality heterozygous variants were selected that (a) had a read depth of at least 10 (b) were not adjacent to a homopolymer exceeding 5 base pairs (c) were present in <5 whole exome sequencing (WES) datasets collected from 147 individuals not affected with HLHS and (d) were present at a frequency <1% (de novo, loss-of-function, CHD panel genes) or <3% (compound heterozygous, hemizygous or homozygous recessive) in the Exome Variant Server (WES data from 6503 individuals, URL: http://evs.gs.washington.edu/EVS) 1000 Genomes (WGS data from 1092 individuals),^3^ and/or Complete Genomics Genome (WGS data from 69 individual).^4^ Second, functional variants were selected, defined as those that impacted a protein sequence, canonical splice site, microRNA coding sequence/binding site, enhancer region, or transcription factor binding site within a promoter validated by ENCODE chromatin immunoprecipitation experiments.^5^ Third, using parental and sibling WGS data, rare, functional variants in the proband were then filtered for those that arose *de novo* or fit homozygous recessive, compound heterozygous, or X-linked recessive modes of inheritance. In addition, any inherited frameshift and start/stop codon variants were retained if they occurred in a gene intolerant to loss-of-function (pLI score > 0.75).

#### WGS of an HLHS cohort and unaffected controls

WGS of 130 unrelated individuals with HLH (80% HLHS, 20% congenital heart disease (CHD) with left ventricular hypoplasia) was performed utilizing the Mayo Clinic Medical Genome Facility. For the control population, 861 individuals from the Mayo Clinic Center for Individualized Medicine’s Biobank repository^6^ were selected based upon absence of personal or family history of CHD and underwent WGS at HudsonAlpha Institute for Biotechnology. Variant call annotation for all 991 individuals was performed by the Mayo Clinic Bioinformatics Core. Whole genome libraries were prepared for 130 individuals with HLHS and 101 bp or 150 bp paired-end sequencing was performed on either the Illumina HiSeq 2000 (n=56) or HiSeq 4000 (n=74), respectively. For the 861 Biobank individuals, whole genome libraries were prepared and 150 base pair paired-end sequencing was carried out on the HiSeqX Ten platform. Reads from all 991 individuals were aligned to the hg38 reference genome using BWA-MEM and duplicate reads were marked using Picard. Local realignment of insertion/deletions (INDELs) and base quality score recalibration was then performed using the Genome Analysis Toolkit version 3.4 (GATK) followed by SNV/INDEL calling with Haplotype Caller and Genotype GVCFs. VerifyBAMID ^7^ was used to estimate sample contamination. Samples with low coverage (<90% of genome covered at 10X) or a high contamination estimate (FREEMIX > 0.03) were excluded. A single VCF file with SNV and INDEL calls from all 991 individuals was created for subsequent statistical analysis.

#### Rare variant burden analysis of *LRP2* and *APOB*

WGS data from cases and controls was compared for rare variant burden of the candidate genes that have been functionally validated in both systems *(LRP2, APOB)* (Supplementary Table **6**). Genotypes with genotype quality (GQ) <20 were excluded, and the resulting data was used to calculate variant call rates and Hardy-Weinberg Equilibrium (HWE) p-values. Variants with call rate <0.95 or HWE p-value <1e-8 were excluded. In addition, variants were required to pass VQSR^1,2,8^. Variants were only included in the analysis if they had a strong predicted functional impact based on annotation information from Clinical Annotation of Variants (CAVA)^9^. Specifically, we included frameshift, nonsynonymous, stop-gain, and stop-loss variants, as well as variants that alter an essential splice site. We further restricted the nonsynonymous variants to include only those with Combined Annotation Dependent Depletion (CADD) scores >24^10^. Rare variants (MAF <0.01 across all races) were identified based on allele frequencies in ExAC, gnomAD, and ESP (WES data from 6503 individuals, URL: http://evs.gs.washington.edu/EVS)^11^. The gene-level, case-control association analysis was conducted using SKAT-O.^12^ Variants were weighted using the beta(1,25) density function of the observed MAF (the default option in SKAT) and were mapped to genes using HG38 gene coordinates from Ensembl^13^. Correcting for multiple testing, the threshold for statistical significance was set at p<0.025 (0.05/2 genes).

After enrichment of rare, predicted-damaging missense variants in *LRP2* was established, we accounted for the potential influence of race and also relaxed functional constraints. Subsequent analyses were confined to 117 individuals with HLHS possessing >80% of ancestral Caucasian alleles. All variants residing in the gene body of *LRP2* were included, in addition to 1000 base pairs upstream of the transcription start site. Variants were isolated and annotated in CAVA utilizing the canonical transcript of *LRP2* (ENST00000263816). In addition to analyzing the total number of variants spanning the gene body, SKAT-O analysis was performed separately for each type of variant in the following categories: missense, intronic, splice site region, splice region (in-frame, missense, synonymous), synonymous, 3’ untranslated region, 5’ untranslated region and 1000 base pairs upstream of the transcription start site. Independent of CAVA annotation, SKAT-O analysis was also performed on regulatory regions as determined by ChIP-Seq data from two different sources. The first analysis included variants within regions of *LRP2* impacted by histone modification and CTCF binding from publicly available ENCODE datasets.^14^ Twenty-one human cardiovascular tissues were assessed prior to confining the analysis to fetal human heart tissue (n=3) (Supplementary Table **9**). The second analysis was confined to ENCODE ChIP-Seq data for 161 transcription factors in 91 cell types (wgEncodeRegTfbsClusteredV3 table in UCSC) (http://genome.ucsc.edu/).

#### iPSC production and spontaneous differentiation of proband/parent cells

Fibroblasts were extracted from tissue by migration onto culture plates in fibroblast medium (DMEM, 10% Fetal bovine serum (FBS), penicillin/streptomycin (P/S), all from Thermo Fisher, Waltham, MA). For the reprogramming process, 5×10^4^ fibroblasts were plated and incubated overnight to allow attachment as previously described^15^. On the infection day, medium was supplemented with lentivirus encoding reprogramming genes *SOX2, OCT4, KLF4* and *c-MYC* and incubated for 12 h. Cells were grown in fibroblast medium for 3 days prior to being passaged onto a matrigel coated plate. Once cells were attached, fibroblast medium was substituted by pluripotency-sustaining medium supplemented with 10μM of ROCK inhibitor (TOCRIS, Bio-Techne, Minneapolis, MN) and refreshed daily until colonies appeared (3-6 weeks). Individual colonies were manually picked and expanded on matrigel coated plates in mTeSR1 medium (STEMCELL Technologies, Vancouver, CA). Approximately every 5-6 days cells were mechanically passaged onto fresh matrigel coated plates.

For spontaneous differentiation cells were treated with collagenase IV (Invitrogen, ThermoFisher) for 20 min, gently dislodged from the plate and transferred into suspension culture in ultra-low attachment 6-well plates in differentiation medium (DMEM/F12, 20% FBS, 1% glutamax, 1% non-essential amino acids, and 0.1% 2-mercaptoethanol). On day 5, floating aggregates were transferred to gelatin-coated tissue culture plates where medium was refreshed every 2 to 3 days. Cells were harvested for RNA extraction on day 25.

#### iPSC characterization of proband/parent cells

For pluripotency characterization cells were fixed in 4% paraformaldehyde for 10 min, permeabilized with 0.1% triton-X, blocked using Superblock and stained for membrane antigens TRA-1-60 (monoclonal mouse IgM 1:100), SSEA3 (rat 1:100) and transcription factor Nanog (rabbit 1:100) (all from Stemgent, Cambridge, MA). Characterization of sarcomeric proteins included staining for MLC2a (monoclonal mouse IgG 1:200, Synaptic Systems, Göttingen, Germany) and MLC2v (rabbit 1:200, Proteintech, Rosemont, IL). Secondary staining consisted of Alexa fluor 568 anti-mouse IgM or IgG, Alexa fluor 488 anti-rat and Alexa fluor 633 or 488 antirabbit, all used at 1:250 dilution (Molecular Probes, Thermo Fisher).^15^. Nuclei were stained with 4’,6-diamidino-2-phenylindole (DAPI). Confocal images were acquired with a Zeiss LSM 510.

Pluripotency properties were determined *in vivo* using a teratoma assay. All studies including animals were approved by the Institutional Animal Care and Use Committee at Mayo Clinic. Half a million cells in 50 μl of a 1:1 solution of differentiation medium and matrigel were injected subcutaneously in each flank of an immunodeficient mouse. Tumor growth was monitored for up to 10 weeks with growing masses harvested as they reached a 1cm^3^ volume. Tissue was flash frozen, cryosectioned and stained using hematoxylin/eosin^15^.

Electron microscopy images were acquired with a JEOL 1200 EXII transmission electron microscope. Cells were processed through fixation with 1% glutaraldehyde and 4% formaldehyde in 0.1 M phosphate buffered saline (pH 7.2), staining with lead citrate and ultramicrotome sectioning prior to imaging^16^.

#### Transcriptome profiling with RNA sequencing (RNA-seq) and bioinformatics analysis

RNA was extracted from iPSCs and differentiated cells at days 0 and 25 using a combination of Trizol and Qiagen RNeasy mini kit columns. Sequencing library was prepared using TruSeq RNA Library Preparation Kit v2. All samples were sequenced on Illumina Hiseq 2000 at Mayo Clinic Medical Genome Facility. The following RNA-seq data analysis was performed on Dell Precision T7500 workstation which has 96 GB RAM and 20 Intel Xeon X5680 processors (3.33GHz) and runs 64-bit Red Hat Enterprise Linux 6.3 (Kernel Linux 2.6.32-279.14.1.el6.x86_64). The alignment of short reads (50 bp) from FASTQ files was performed using Bowtie2 and Tophat2 software. All mapped reads were assembled to transcripts using Cufflinks and merged together using Cuffmerge. Differential analyses were performed between proband and parents at each time point (day 0 and day 25). The results from Cuffdiff were imported into a SQL database using R package CummeRbund for extracting significantly differential genes and other data manipulation. Differential genes were selected based on the default setting in Cuffdiff with adjusted p values at 0.05 after FDR control for correcting multiple hypothesis tests and a minimum fold change of ± fold or greater relative to control lines. Bioinformatics analysis of gene expression changes was performed using available online tools to describe differential patterns between proband, mother and father. Gene functional annotation and classification was generated using the Database for Annotation, Visualization and Integrated Discovery bioinformatics module (http://david.abcc.ncifcrf.gov). Additionally, mapping was performed using the Kyoto Encyclopedia of Genes and Genomes array tool (http://www.kegg.jp/kegg/download/kegtools.html). Heat maps were generated from sorted Database for Annotation, Visualization and Integrated Discovery and Kyoto Encyclopedia of Genes and Genomes gene subsets using TIGR’s open source MeV software (http://tm4.org/mev). In each sample, for each mapped gene, sample data points were normalized to the mean expression across proband, father and mother and subsequently log2 transformed. Significant function groups were ranked based on statistical significance (p) from hypergeometric distribution.

#### Guided cardiac differentiation

Guided differentiation was achieved using a modified version of a previously published protocol.^17^ In brief, iPSCs cells were cultured as monolayer for two passages prior to induction. Next, they were treated with 8-12 μM Wnt activator CHIR99021 (Stemcell technologies) for 20 h followed by 24 h wash out period in DMEM:F12 with B27 supplement (Gibco, ThermoFisher). Medium was then refreshed and supplemented with 5 μM Wnt inhibitor IWP2 for 48 h. Cells were maintained in DMEM:F12 plus B27 for an extra 48 h and in DMEM:F12 plus B27 (minus insulin) thereafter. Cultures were sampled for RNA extraction before induction as well as at days 1, 3, 5, 7, 14 and 37. Beating could be observed after 7-10 days of differentiation.

#### hiPSCs, hiPSC-CMs, siRNA transfection, EdU assay, Immunostaining and qRT-PCR

hiPSCs derived from HLHS families were plated in 384 wells coated with matrigel at 5000 cell/well density using mTeSR-1 (Stem Cell). After 3 days, EdU was added to the media and was incubated for 1 hour. Cells were fixed in 4% PFA and stained for EdU and DAPI (Invitrogen). EdU was detected using Click-it Plus EdU Imaging Kit (Life Technologies) following manufacturing directions. iPSC-CMs from 5H family and generic hiPSC-CMs at day 25 of differentiation were plated in 384 wells coated with matrigel at 5000 cells/well density in Maintenance Media (MM) (RPMI, 2% KOSR, 1% B27, 1% P/S). siRNA transfection was performed using Opti-Mem (Gibco) and Lipofectamine RNAiMAX (Gibco). siRNAs were purchased at Dharmacon and used at a final concentration of 25 nM. siRNA transfection efficiency was tested with qRT-PCR (Supplementary Figure **2C**). siRNA scramble was used as control (siCTR). To test WNT pathway interaction, cells were treated with 1 uM BIO (GSK-3 inhibitor) (Sigma B1686) for three days. Two days after transfection, 50% of media was removed and replaced with 20 μM EdU in MM media. 24 hours after, cells were fixed in 4% paraformaldehyde (PFA) and blocked in blocking buffer (10% Horse Serum, 10% Gelatin, 0.5% Triton X-100. Cells were stained with mouse monoclonal anti-α-Actinin (ACTN1) (Sigma, A7811 1:800), secondary antibody Alexa Fluor 568 anti-mouse (Invitrogen, 1:1000) and DAPI (1:1000) in blocking buffer and imaged using ImageXPress microscope, (Molecular Devices) and analyzed with MetaXpress Analysis software (Molecular Devices). To obtain a cardiomyocyte proliferation index, the total number of cells positive for EdU, α-Actinin and DAPI was divided by the total number of DAPI cells and expressed as percentage. For qRT-PCR experiments, total RNA was extracted using TRIzol and chloroform. 1ug of RNA was converted in cDNA using QuantiTect Reverse Transcription kit (Qiagen). qRT-PCR was performed using Syber green (Biorad). Human primers sequences for qRT-PCR were obtained from Harvard Primer Bank. TP53 (Primer Bank ID: 371502118c1), CDKN1A (Primer Bank ID: 310832423c1), LRP2 (Primer Bank ID: 126012572c1), FZD10 (Primer Bank ID: 314122154c3), PTCH1 (Primer Bank ID: 134254431c3), CCNE1 (Primer Bank ID: 339275820c3) PCNA (Primer Bank ID: 33239449c1), CCNB1 (Primer Bank ID: 356582356c1), CCNB2 (Primer Bank ID: 332205979c1), CDK1 (Primer Bank ID: 281427275c1), CRADD (Primer Bank ID: 51988883c1), CASP6 (Primer Bank ID: 73622127c1), CDKN2C (Primer Bank ID: 17981697c1), CDKN1C (Primer Bank ID: 169790898c1), APOB (Primer Bank ID: 105990531c1), ELF4 (Primer Bank ID: 187608766c1), HN1(JPT1) (Primer Bank ID: 7705877a1), HS6ST2 (Primer Bank ID: 116295253c2), HSPG2 (Primer Bank ID: 140972288c1), PRTG (Primer Bank ID: 224500891c2), SDHD (Primer Bank ID: 222352156c3), SIK1 (Primer Bank ID: 116256470c1), SLC9A1 (Primer Bank ID: 381214343c3). GAPDH (Primer Bank ID: 378404907c1) was used as housekeeping gene and used to normalize the data. At least 3 independent biological replicates were performed for each experiment.

#### Quantitative Real Time PCR (qRT-PCR) in iPSC

qRT-PCR for pluripotency and disease associated markers was performed in iPSC samples. RNA was extracted using a combination of Trizol and Qiagen RNeasy mini kit columns. cDNA for pluripotency assessment was synthesized using reverse transcriptase supermix reagents (Invitrogen, Thermo Fisher). In the case of expression levels during a time course of differentiation, a Biorad (Hercules, CA) iScript synthesis kit was used. qRT-PCR was performed using pre-designed primers (IDT Integrated DNA technologies, Coralville, IA) for *CDH1* (Hs00170423_m1), *DNMT3B* (Hs01003405_m1), *DPPA2* (Hs00414521_g1), *DPPA5* (Hs00988349_g1), *ERAS* (Hs.PT.45.4849266.g), *GDF3* (Hs00220998_m1), *OCT4* (Hs.PT.45.14904310.g), *REXO1* (Hs.PT.45.923095.g), *SALL4* (Hs00360675_m1), *TDG1* (Hs02339499_g1) and *TERT* (Hs99999022_m1) in the characterization of the pluripotent state and *APOB* (Hs.PT.56a.1973344), *DHCR24* (Hs.PT.56a.4561516), *ELF4* (Hs.PT.56a.25941471), *HN1* (Hs.PT.58.40922463.g), *HSPG2* (Hs.PT.56a.18698732), *HS6ST2* (Hs.PT.56a.1354985), *LRP2* (Hs.PT.56a.1584067), *MYLK* (Hs.PT.56a.39795491), *PCDH11X* (Hs.PT.56a.26531358), *PRTG* (custom design), *SIK1* (Hs.PT.58.2995158), *SLC9A1* (Hs.PT.58.15072523), and *SDHD* (Hs.PT.58.40267655.g) for expression during guided cardiac differentiation. All values were normalized to *GAPDH* (Hs.PT.45.832633).

#### Statistical analysis

The qPCR time course gene expression data were analyzed using Generalized Linear Model (GLM) to assess the statistical significance. EdU incorporation experiments and pTP53 staining were analyzed with GraphPad Prism 8. For both, *p*<0.05 was considered significant.

## Zebrafish Husbandry

All zebrafish experiments were performed in accordance to protocols approved by IACUC. Zebrafish were maintained under standard laboratory conditions at 28.5°C. In addition to Oregon AB wild-type, the following transgenic lines were used: *Tg(myl7:EGFP)^twu277^^1^* and *Tg(myl7:H2A-mCherry)^sd12^*^2^.

## Zebrafish SOHA (Semi-automated Optical Heartbeat Analysis)

Larval zebrafish (72 hpf) were immobilized in a small amount of low melt agarose (1.5%) and submerged in conditioned water. Beating hearts were imaged with direct immersion optics and a digital high-speed camera (up to 200 frame/sec, Hamamatsu Orca Flash) to record 30 seconds movies; images were captured using HC Image (Hamamatsu Corp.). Cardiac function was analyzed from these high-speed movies using semi-automatic optical heartbeat analysis software^3,4^, which for zebrafish quantifies heart period (R-R interval), cardiac rhythmicity, as well as chamber size and fractional area change. All hearts were imaged at room temperature (20-21·C). Statistical analyses were performed using Prism software (Graphpad). Significance was determined using two-tailed, unpaired student t-test or one-way ANOVA and Dunnett’s multiple comparisons post hoc test as appropriate.

## Zebrafish Cardiomyocyte Cell Counts and Cardiac immunofluorescent imaging

To count cardiomyocytes, we used the expression of H2AmCherry in the nuclei (*Tg(myl7:H2A-mCherry)^2^* to qualify as an individual cell, performed the “Spot” function in Imaris to distinguish individual cells in reconstructions of confocal z-stacks^5,6^. To compare data sets, we used Prism software (GraphPad) to perform Student’s t-test with two-tail distribution. Graphs display mean and standard deviation for each data set.

Whole-mount immunofluorescence was performed as previously described^5–7^, using primary monoclonal antibodies against the following, and secondary antibody, either donkey anti-chicken AlexaFluor 488 or donkey anti-rabbit AlexaFluor 568, was used in 1:200 dilution:

| Primary Antibody | Host/Reaction | Company | Part# | Concentration |
| --- | --- | --- | --- | --- |
| GFP | Chicken | Aves Labs | GFP-1020 | 1:300 |
| mCherry | Rabbit | Rockland | 600-401-P16S | 1:200 |
| **Secondary Antibody/Stains** |  |  |  |  |
| AlexaFluor 488 | Donkey anti Chicken | Jackson ImmunoResearch | 703-545-155 | 1:200 |
| AlexaFluor 568 | Donkey anti Rabbit | Invitrogen |  | 1:200 |
| DAPI (500mg/ml) |  | Invitrogen |  | 1:200 |

Confocal imaging was performed on an LSM 710 confocal microscope (Zeiss, Germany) with a 40x water objective. Exported z-stacks were processed with Imaris software (Bitplane), Zeiss Zen, and Adobe Creative Suite software (Photoshop and Illustrator 2020). All confocal images shown are projection views of partial reconstructions from multiple z-stack slices, except where noted that images are views of a single slice.

## Zebrafish CRISPR/Cas9 experiments

Detailed steps for *lrp2a* were previously described^8^ and we followed IDT manufacture instruction for Complexes preparation. <u>crRNA:tracrRNA Duplex Preparation</u>: Target-specific Alt-R^®^ crRNA (Dr.Cas9.LRP2A.1.AC, /AltR1/rCrC rCrUrC rGrCrU rUrArU rArUrU rCrUrC rCrArA rGrUrU rUrUrA rGrArG rCrUrA rUrGrC rU/AltR2/) and common Alt-R^®^ tracrRNA were synthesized by IDT and each RNA was dissolved in duplex buffer (IDT) as 100 μM stock solution. Stock solutions were stored at −20°C. To prepare the crRNA:tracrRNA duplex, equal volumes of 100 μM Alt-R^®^ crRNA and 100μM Alt-R^®^ tracrRNA stock solutions were mixed together and annealed by heating followed by gradual cooling to room temperature by manufacture instruction: 95°C, 5 min on PCR machine; cool to 25°C; cool to 4°C rapidly on ice. The 50 μM crRNA:tracrRNA duplex stock solution was stored at −20°C. <u>Preparation of crRNA:tracrRNA:Cas9 RNP Complexes</u>: Cas9 protein (Alt-R^®^ S.p. Cas9 nuclease, v.3, IDT) was adjusted to 25 μM stock solution in 20mM HEPES-NaOH (pH 7.5), 350 mM KCl, 20% glycerol, dispensed as 8 ul aliquots, and stored at – 80°C. 25μM crRNA:tracrRNA duplex was produced by mixing equal volumes of 50 μM crRNA:tracrRNA duplex stock and duplex buffer (IDT). We used 5 μM RNP complex. To generate 5μM crRNA:tracrRNA:Cas9 RNP complexes: 1 μl 25 μM crRNA:tracrRNA duplex was mixed with 1 μl 25 μM Cas9 stock, 2 μl H2O, and 1 μl 0.25% phenol red solution. Prior to microinjection, the RNP complex solution was incubated at 37°C, 5 min and then placed at room temperature. Approximately one nanoliter of 5 μM RNP complex was injected into the cytoplasm of one-cell stage embryos to generate F0 larva.

## SUPPLEMENTAL RESULTS

### Whole Genome Sequencing

WGS was carried out on genomic DNA samples from all five family members, based on 101 base paired-end reads that passed quality control standards; 92% of the reads mapped to the genome. After marking and filtering out duplicate reads, over 99% of the hg19 human reference genome had coverage. The average depth across the genome was 36X and an average of 91% of the gene body regions (exons, introns, and 5’ and 3’ untranslated regions) demonstrated a minimal read depth of 20 reads.

## SUPPLEMENTARY FIGURES

**Supplementary Figure 1:**
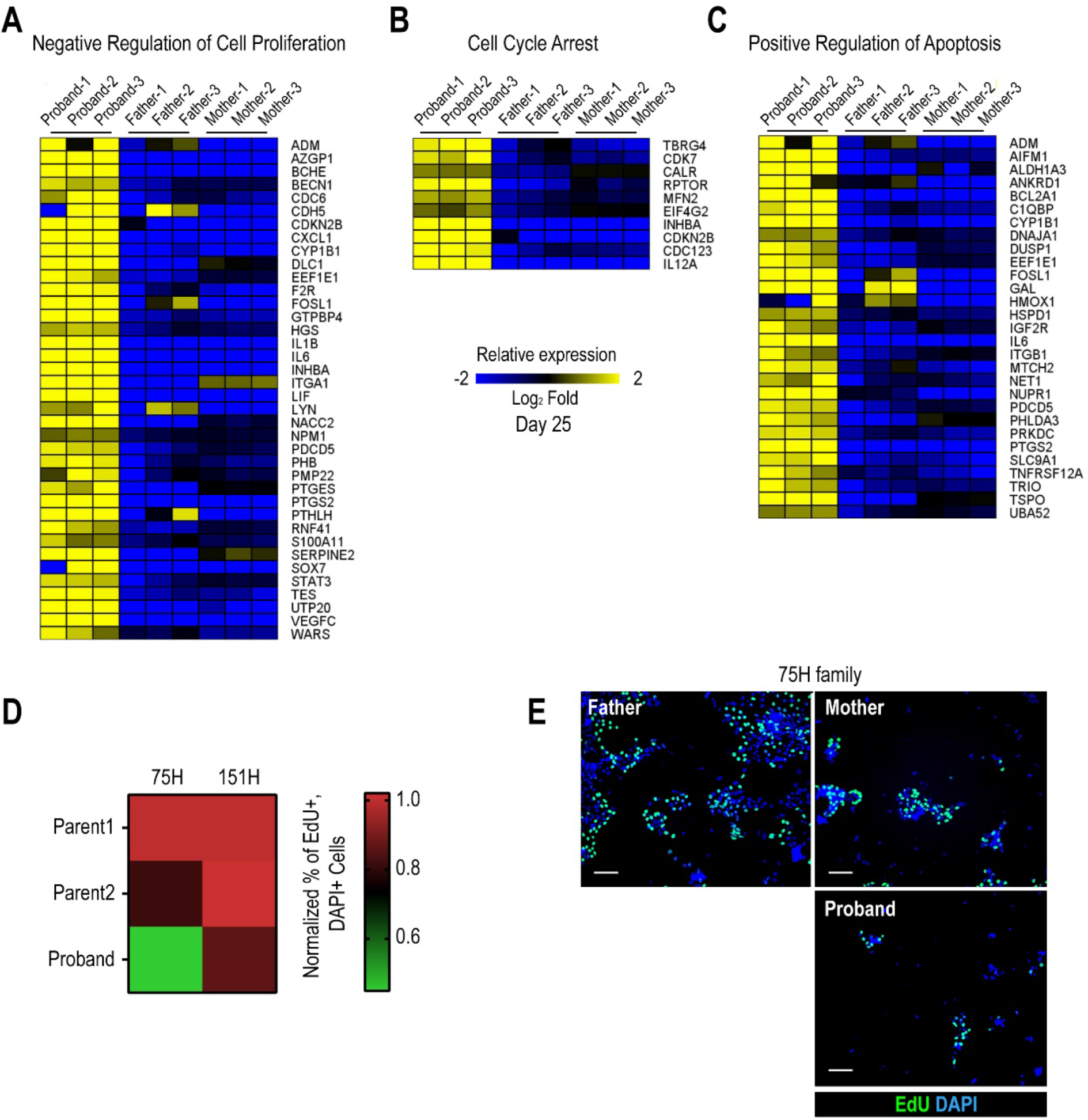
Cell cycle activity is altered in HLHS patient-derived iPSCs and CMs. (**A**) Heatmap of negative regulation of cell proliferation associated genes from RNA-seq experiments in proband vs parents. (**B**) Heatmap of cell cycle arrest associated genes. (**C**) Heatmap of positive regulation of apoptosis associated genes. (**D**) Quantification of percentage of EdU+ cells in 75H and 151H iPSCs. Proband iPSCs show reduced EdU incorporation compared to the father. *p<0.05 for 151 H proband as compared to both parents **p<0.01 for 75H proband as compared to both parents, One-way ANOVA. (**E**) Representative images of 75H family iPSCs labeled for EdU and DAPI. Scale bar: 50 μm.

**Supplementary Figure 2:**
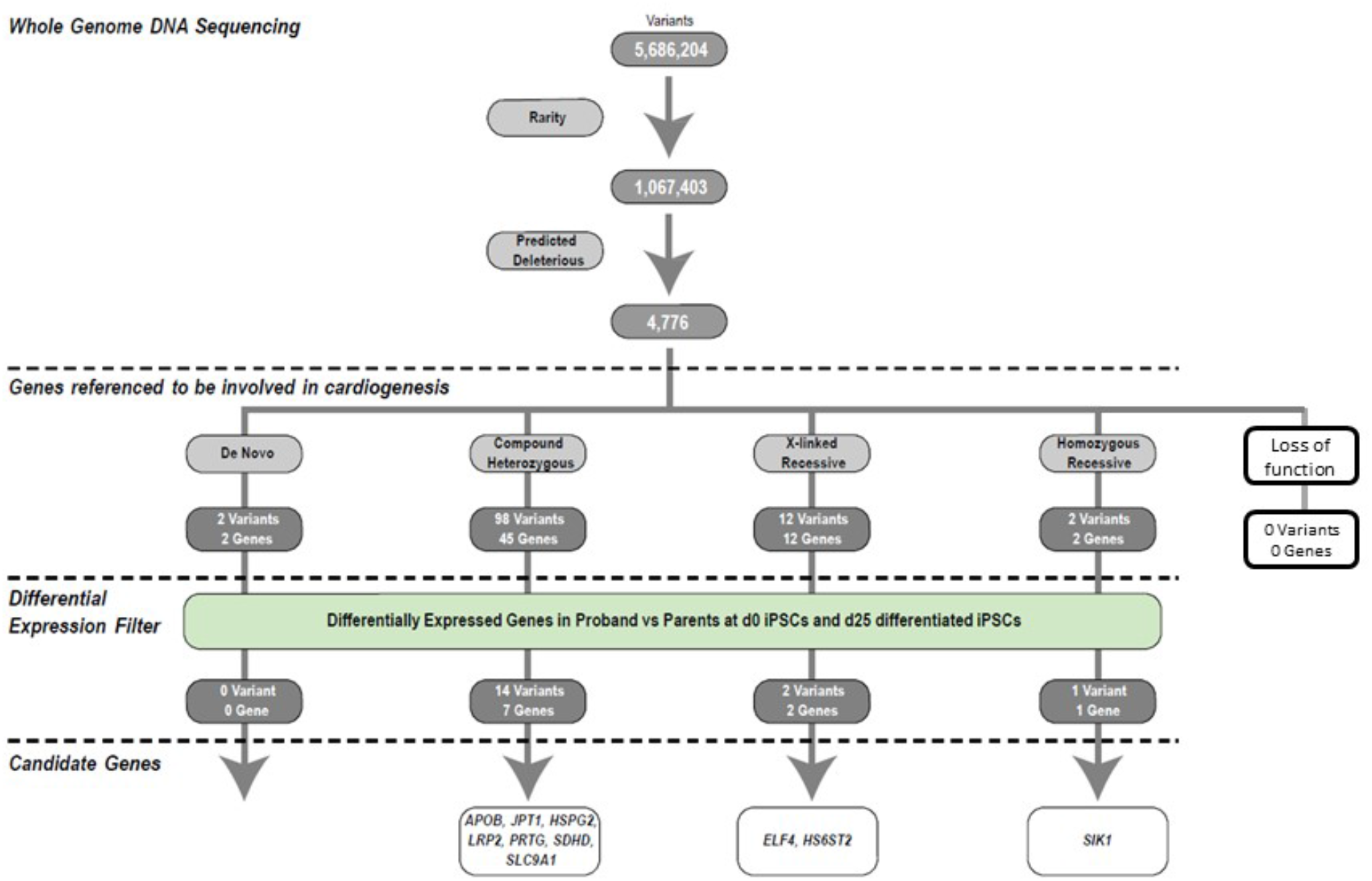
Identification of HLHS candidate genes from whole genome and RNA sequencing. An iterative, family-based variant filtering approach based on rarity, functional impact, and mode of inheritance yielded 61 candidate genes. RNA sequencing data from d0 iPSC and d25 differentiated iPSC were used to filter for transcriptional differences yielding ten candidate genes.

**Supplementary Figure 3:**
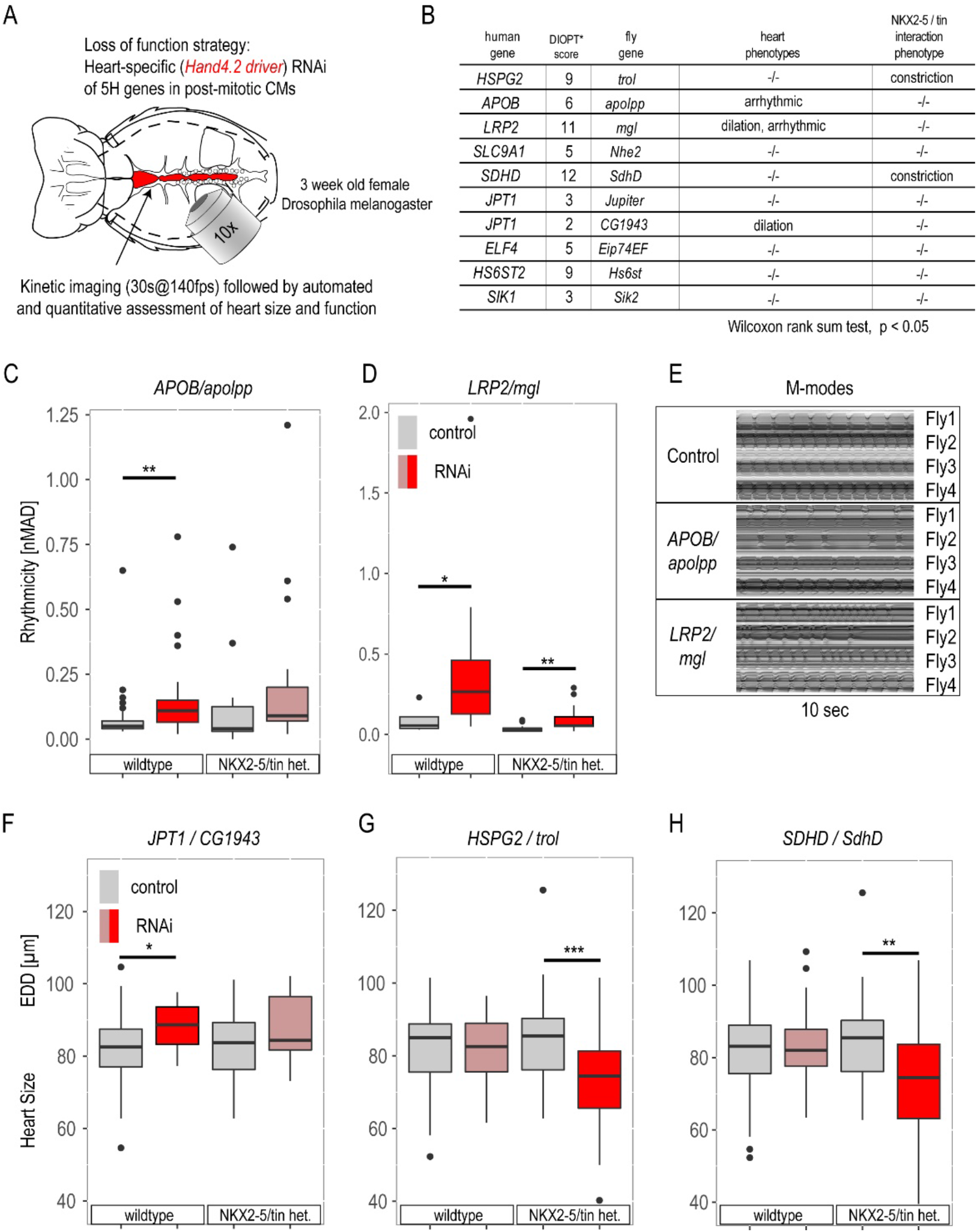
Phenotypic assessment of HLHS candidate genes in *Drosophila* adult hearts. (**A**) Schematic of the *Drosophila* adult heart assay. (**B**) Human candidate genes and corresponding Drosophila ortholog as determined by DIOPT score (*confidence score: number of databases reporting orthology). Listed are heart phenotypes upon knockdown (KD) in wildtype or NKX2-5/tin+/− heterozygous background. (**C-E**) RNAi-induced arrhythmicity and M-modes observed with *LRP2/mgl* and *APOB/apolpp* KD. (**F-H**) Heart size (EDD: end-diastolic diameter) alterations upon RNAi-KD of JPT1, HSPG2 and SDHD (also in NKX2-5/tin heterozygous background). Wilcoxon rank sum test: *p<0.05, **p<0.01, **p<0.001.

**Supplementary Figure 4.**
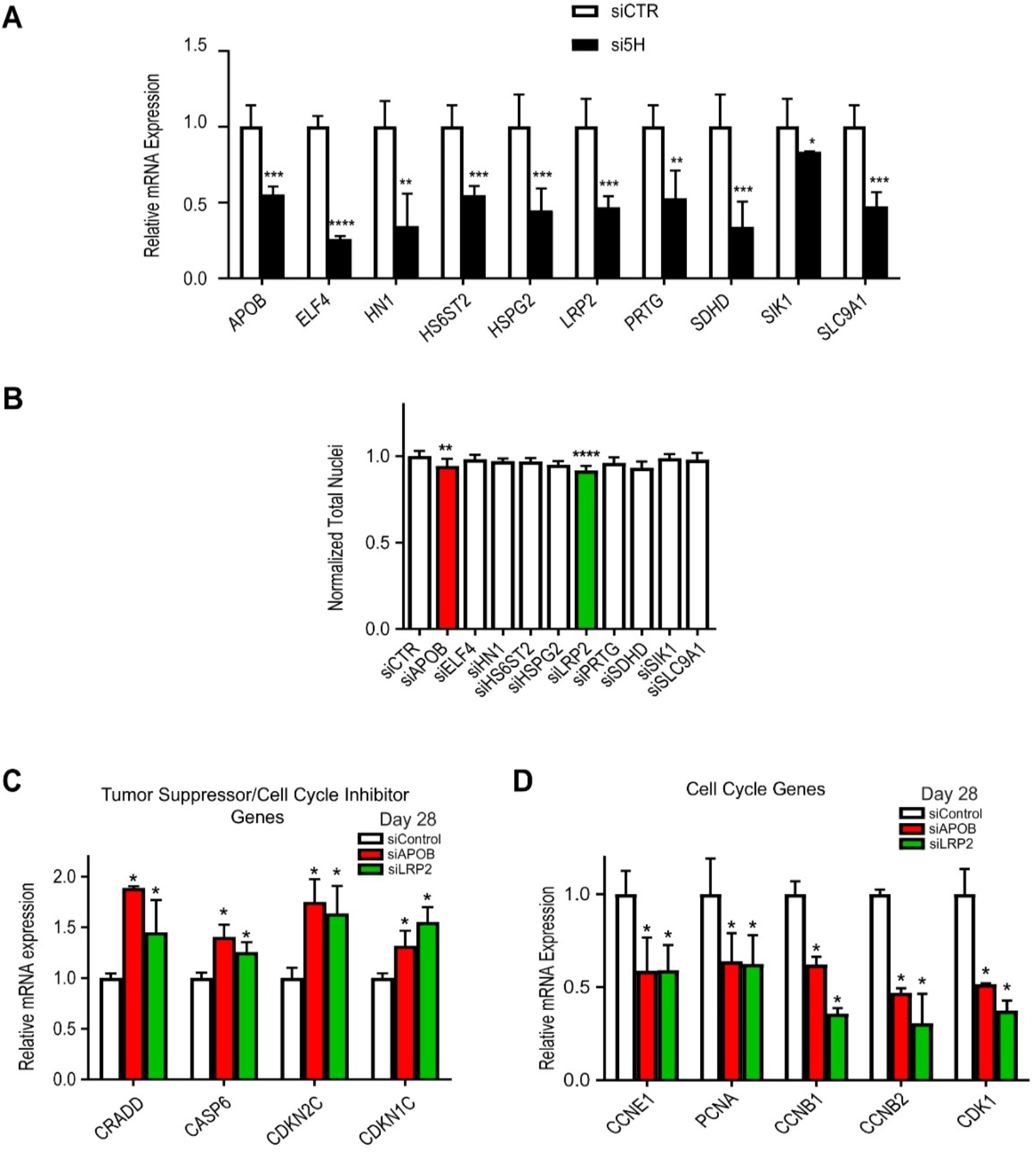
*LRP2* and *APOB* KD reduces total nuclei and affect cell cycle in hiPSC-CMs. (**A**) qPCR showing KD efficiency for the ten candidate genes in day 28 hiPSC-CM. For all the ten genes tested, siRNAs KD efficiency is about 50%. *p<0.05, **p< 0.01, ***p<0.001, ****p< 0.0001, Student’s t-test. (**B**) Quantification of total nuclei in day 28 hiPSC-CMs transfected with siRNAs directed to ten candidate HLHS genes. *LRP2* and *APOB* KD reduce total cells number. (**C**) qPCR of tumor suppressors/cell cycle inhibitor genes in day 28 hiPSC-CMs upon KD of *LRP2* or *APOB.* Tumor suppressors/cell cycle inhibitors are up-regulated upon *LRP2/APOB* KD compared to control. *p<0.05, Student’s t-test. (**D**) qPCR of cell cycle genes in day 28 hiPSC-CMs upon KD of APOB or LRP2. Cell cycle genes are downregulated upon *LRP2/APOB* KD. *p<0.05, Student’s t-test.

**Supplementary Figure 5:**
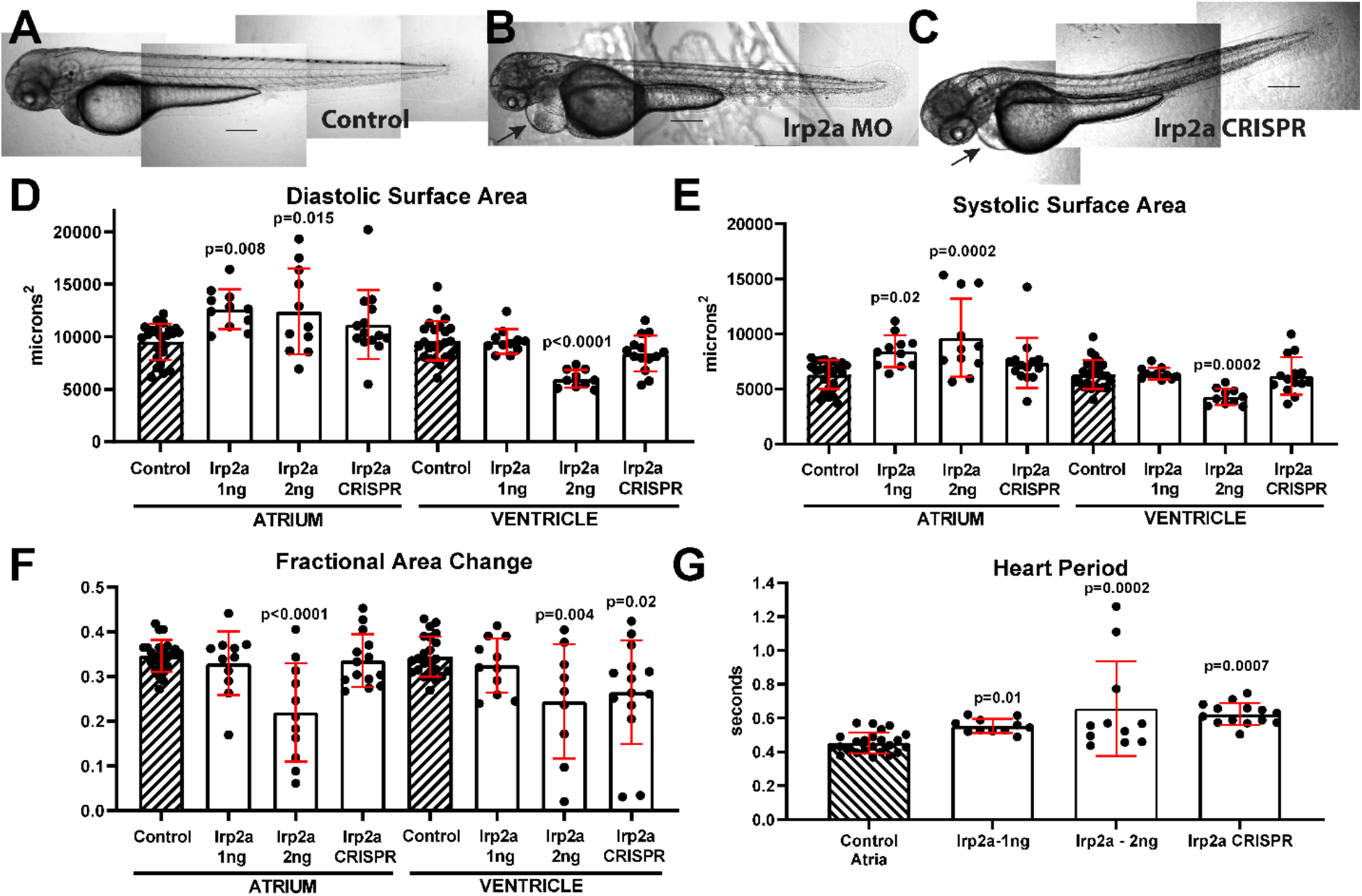
*Irp2a* KD and CRISPR causes reduced contractility and bradycardia in zebrafish larva. **(A)** Uninjected zebrafish larva, **(B)** *lrp2a* morpholinos (MO) (1 pl of 2 ng/μl) injected larva and **(C)** *lrp2a* CRISPR genome edited F0 larva all exhibit relatively normal body morphology at 72 hpf, note the pericardial edema evident in morphant and F0 mutant (arrows). Scale bar: 200 μm. **(D)** End Diastolic surface area and **(E)** End Systolic surface area in atria and ventricles determined from high speed movies of beating hearts at 72 hpf. **(F)** Contractility, measured as fractional area change, was significantly reduced in ventricles from both morphants and mutants. **(G)** Heart period was significantly lengthened in morphants and mutant larva. (For D-G) Two doses of *lrp2a* MO were injected (1 pl of 1 ng/μl or 2 ng/μl). For CRISPR/Cas9 and guide RNA concentration, please see detailed information in the Materials and Methods section. One-way ANOVA, Dunnett’s multiple comparisons post hoc test.

## SUPPLEMENTARY TABLES

**<u>Supplementary Table 1</u>**

Please see separately attached excel file, Supplemental Table 1.

**<u>Supplementary Table 2</u>**

Please see separately attached excel file, Supplemental Table 2.

**<u>Supplementary Table 3</u>**

Please see separately attached excel file, Supplemental Table 3.

**<u>Supplementary Table 4</u>**

Please see separately attached excel file, Supplemental Table 4.

**<u>Supplementary Table 5</u>**

Please see separately attached excel file, Supplemental Table 5.

**<u>Supplementary Table 6</u>**

Please see separately attached excel file, Supplemental Table 6.

**<u>Supplementary Table 7</u>**

Please see separately attached excel file, Supplemental Table 7.

**<u>Supplementary Table 8</u>**

Please see separately attached excel file, Supplemental Table 8.

**<u>Supplementary Table 9</u>**

Please see separately attached excel file, Supplemental Table 9.

## SUPPLEMENTARY MOVIES

**Supplemental Movie Legends S1-S3:** Representative heart movies of dissected adult females showing arrhythmic beating pattern in APOB-RNAi (Movie S2) and *LRP2-RNAi* (Movie S3) compared to control hearts (Movie S1). All movies are imaged at 140 frames/sec.

## Notes

### Competing Interest Statement

The authors have declared no competing interest.

### Summary of Updates

This version contains additional data from zebrafish experiments and small molecule treatment of iPSC cells.

