## Supplemental Table 8 for "Patient-specific functional genomics and disease modeling suggest a role for LRP2 in hypoplastic left heart syndrome"

**Supplemental Table 8. Case-Control Association Analysis of Rare Variants in *LRP2***

| | | HLHS (n=117) | | | Controls (n=861) | | | p_value |
|---|---|---|---|---|---|---|---|---|
| | CAVA | Total SNVs | Total Subjects | | Total SNVs | Total Subjects | | |
| Total number of variants (no predicted-damaging threshold) | All | 1568 | 117 | 100.00% | 10772 | 861 | 100.00% | 0.0035 |
| Missense | NSY | 31 | 29 | 24.80% | 147 | 142 | 16.50% | 0.0178 |
| Splice Region Intron | SS | 2 | 2 | 1.70% | 19 | 19 | 2.20% | 0.2206 |
| Splice Region: Inframe, Missense, Synonymous | EE | 2 | 2 | 1.70% | 16 | 16 | 1.90% | 1 |
| Synonymous | SY | 15 | 14 | 12.00% | 83 | 79 | 9.20% | 0.1545 |
| 1 KB Upstream | N/A | 4 | 4 | 3.40% | 18 | 18 | 2.10% | 0.2976 |
| 5' untranslated region | 5PU | 0 | 0 | 0.00% | 1 | 1 | 0.10% | 0.5567 |
| 3' untranslated region | 3PU | 10 | 7 | 6.00% | 38 | 37 | 4.30% | 0.0934 |
| Intron | INT | 1504 | 117 | 100.00% | 10450 | 861 | 100.00% | 0.0082 |
| Variant lies within an within active histone marks from CHIPSeq data confined to human cardiovascular tissue (n=21) | N/A | 53 | 39 | 33.30% | 374 | 285 | 33.10% | 0.6301 |
| Variant lies within an within active histone marks from CHIPSeq data confined to human fetal heart (n=3) | N/A | 25 | 22 | 18.80% | 158 | 149 | 17.30% | 0.5692 |
| Variant lies within a transcription factor binding site | N/A | 94 | 60 | 51.30% | 590 | 394 | 45.80% | 0.1847 |
